# Chemotaxis overrides Barotaxis during Directional Decision-Making in *Dictyostelium discoideum*

**DOI:** 10.1101/2020.01.14.904748

**Authors:** Yuri Belotti, David McGloin, Cornelis J. Weijer

## Abstract

Neutrophils and dendritic cells have, besides their well characterised chemotactic movement responses, been shown to be able to detect and respond to local differences in hydraulic resistance (*barotaxis*). Furthermore, for neutrophils, it has been suggested that barotaxis overrides chemotaxis. Here, we investigate whether Dictyostelium cells also respond to hydraulic resistance or primarily to chemical gradients using an asymmetric bifurcating micro-channel. This channel design allows us to decouple hydraulic and chemical stimuli, by providing a choice between moving up a chemical gradient or down a chemical gradient into a channel with 100 times lower hydraulic resistance. Under these conditions chemotaxis always overrides barotaxis. Cells confronted by a microchannel bifurcation are observed to often partially split their leading edge and to start moving into both channels. Cells in steeper cAMP gradients, that move faster, split more readily. The decision to retract the pseudopod moving away from the cAMP source is made when the average velocity of the pseudopod moving up the cAMP gradient is 20% higher than the average velocity of the pseudopod moving down the gradient. Surprisingly, this decision threshold is independent of the steepness of the cAMP gradient and speed of movement. It indicates that a critical force imbalance threshold underlies the repolarisation decision.

**Significance Statement:** We investigate the directional ‘decision-making’ of Dictyostelium discoideum cells migrating within engineered micro-channels harbouring asymmetric bifurcations. Unlike neutrophils and immature dendritic cells, Dictyostelium cells strongly prioritise chemical over barotactic guidance cues. Cells in steeper cAMP gradients migrate at higher speeds, split their leading edges more readily when confronted with a bifurcation in the channel. The decision to retract a pseudopod pointing in an unfavourable direction occurs when a critical tension gradient between two competing pseudopods is surpassed. These experiments show that although barotaxis is not a major guidance cue, cellular mechanics plays a major role in leading edge dynamics, including front splitting and polarisation and retraction.

## Introduction

Cell migration plays a key role in several different biological processes, such as embryonic morphogenesis, immune responses and wound healing (1, 2). Various animal cells exhibit extensive migratory capabilities, for instance macrophages and neutrophils crawl towards invaders and engulf and destroy them, osteoclast and osteoblast ensure the continuous remodelling of bones, fibroblast migrate to damaged sites of tissue helping to rebuild them (3). Cell movement is also a key driver of some pathological processes such as osteoporosis, chronic inflammatory diseases and tumour metastasis (1). Insights into the mechanisms that control and execute migration will be required for more effective medical treatments and facilitate new approaches in regenerative medicine and tissue engineering.

One of the most important questions in understanding cell movement is how the cell interprets external cues and actuates the internal cytoskeletal machinery to achieve the motion (4). A variety of biochemical and physical cues have been shown to trigger cellular responses (5–11). Chemical concentration gradients are one of the environmental signals, which can instruct the migration of certain cell types. This kind of response is known as *chemotaxis* and involves a directed migration as a consequence of directional sensing and has been extensively investigated in *in-vitro* systems (12). However, in their physiological environment, cells are exposed to a combination of a variety of chemical and mechanical stimuli and it is still largely unresolved how responses are prioritised and coordinated. The advent of microfluidic techniques has enabled investigation of cell migration in more detail by providing a better control over the mechanical and chemical complexity of the microenvironment that surrounds each individual cell (13). Microfluidics provides good control over dynamics of signalling, as well as over spatial complexity of the cellular environment. For instance, maze-like microfluidic networks (14) have been developed to analyse the mechanisms that amoeboid cells such as neutrophils and Dictyostelium discoideum cells use to effectively navigate through these complex environments (15, 16).

*Dictyostelium discoideum* (Dd) is a well-established model for the study of eukaryotic chemotaxis (17, 18). Research conducted on the mechanism of chemotaxis in this organism has greatly contributed to our basic understanding of chemotaxis and also led to the establishment of novel experimental methods to study chemotaxis now successfully used in other systems (19–24). The signal transduction pathways involved in chemical gradient sensing and transduction to the organisation of the actin myosin cytoskeleton resulting in directed motion are highly conserved between Dictyostelium and neutrophils reviewed in (25). Most of the knowledge on the mechanisms governing cell migration arose from investigations on planar surfaces, but recent studies showed that this classical picture of cell locomotion is inadequate to recapitulate the properties of cell migration within tissues and other complex environments (26).

During *in-vivo* cell migration, a number of physical parameters such as mechanical properties, geometry, adhesion, degree of confinement imposed by the micro-environment in which cells move in, affect how cells respond to chemical cues (27–29). Interestingly, Prentice-Mott *et al.* (30, 31) first showed that Human promyelocytic leukaemia cells, also known as HL60 cells, can interpret small differences in hydraulic resistance when confronted with asymmetric bifurcating channels of varying resistance and called this ‘*barotaxis*’. The authors developed a microfluidic-based migration assay where HL60 cells were forced to move through asymmetric microchannels and observed that cells migrated towards the path of least hydraulic resistance, both in the presence and in the absence of a chemoattractant, showing that the mechanical stimulus seemed to override the chemotactic response.

Here we design and use a novel microfluidic device harbouring microchannels that allow us to decouple the chemical and hydraulic stimuli. The particular topology consists of bifurcating microchannels where only one arm is connected to the source of chemoattractant and has hundred times higher hydraulic resistance with respect to the other arm. In this way cells are subjected to a dual choice: one direction corresponds to an increasing chemical gradient, the other to a significantly lower hydraulic resistance. This approach showed that cells always migrate up the chemical gradient despite the hundred times higher hydraulic resistance and chemotaxis thus overrides barotaxis in Dictyostelium. We furthermore analyse the dynamics of splitting of the leading edge when the cells encounter the bifurcation in the channel as well as the dynamics of retraction of pseudopods pointing in an unfavourable direction. We identify the existence of a response threshold that is independent of the cAMP concentration gradient but dependent on the tension gradient between the competing pseudopods.

## Results

First, we sought to determine whether Dictyostelium cells could detect small pressure differences and show a barotactic response. We constructed microfluidic chips with bifurcating channels of different geometries (Figure 1A-C). In these experiments, cells are enticed to move into thin tight-fitting channels under control of an imposed chemotactic gradient. Initially, diffusion sets up a cAMP gradient from the cAMP loading channel to the cell loading channel (Fig. S1A). Under our experimental conditions the cells detect the chemoattractant as soon as minimal amounts start to diffuse out in the cell loading channel and cells move rapidly in the narrow migration channel towards the source of cAMPS (movie S1). While doing so the cells effectively plug the microfluidic channels. Since diffusion of small molecules is fast on small length scales, the concentration of chemo-attractant at the front and at the back of the cell migrating in the narrow migration channel rapidly equilibrates with the concentrations in the cAMP loading and cell loading channels, respectively. The front of the cell sees the cAMP concentration of the chemo-attractant loading channel (20, 100 or 100nM) and the back the ambient cAMP concentration of the cell loading compartment (close to 0 M cAMP). This results in a steep chemo-attractant gradient over the length of the cell (Fig. S1B, movie S2).

**Figure 1.**
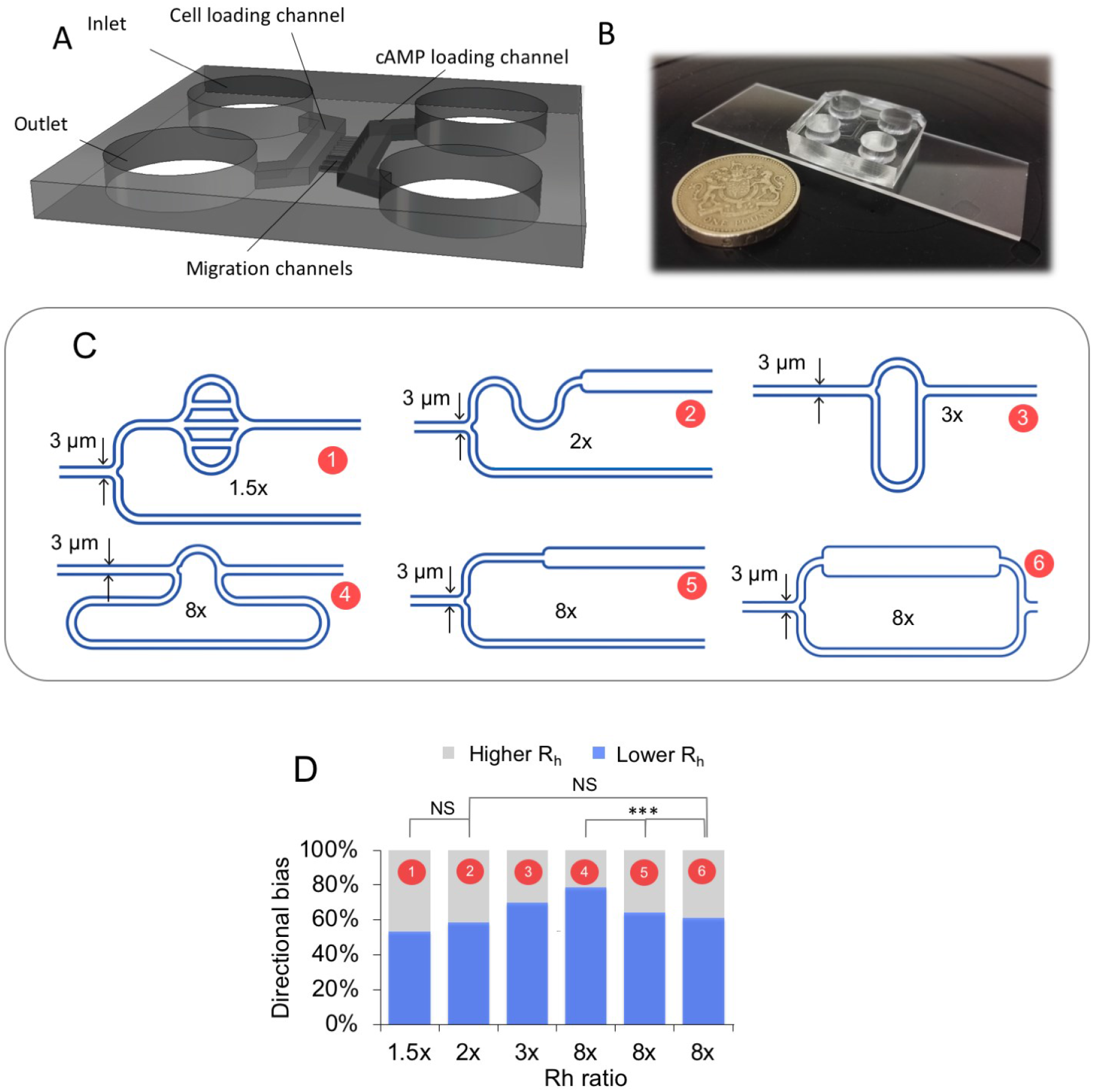
Investigation of barotactic response. (A) 3D structure of the chip, the two symmetric sides are arranged in a ‘ladder-like’ structure. Each side comprises an inlet and an outlet, connected by a loading channel. The two symmetric halves are bridged by a number of small diameter migration channels. (B) Picture of an actual PDMS chip compared to a one pound coin. (C) Schematics of the various geometries of the migration channels used to test the role of barotaxis in the directional bias. The length and width of the channels are designed following the definition of hydraulic resistance of a duct (Eq. 2), whereas the height is set to 2.5 μm. For each design the upper arm corresponds to the arm of least hydraulic resistance. For each geometry, the ratio between the higher and lower resistance is stated. (D) The directional bias, *i.e.* the percentage of cells migrating towards the arm of least hydraulic resistance as a function of the ratio in hydraulic resistance between the two bifurcating arms, for each of the designs shown in (C). To test the statistical significance of any difference in cell behaviour in different geometries Fisher exact test (2×2 and 2×3) was used. The cases 2×, and 8× (geometry 2 and 6) are not statistically different, 2×2 Fisher exact test, p = 0.99, while there is a significant difference in the choice among the different geometries displaying an 8x fold ratio in hydraulic resistance (geometry 4, 5 and 6), 2×3 Fisher exact test, p = 0.00125)

When cells migrate through the asymmetric bifurcating microchannels, they tend to protrude two pseudopods, one in each arm of the bifurcation and after a ‘tug-of-war’ one pseudopod ‘wins’ while the other starts retracting, similarly to what has been reported by Prentice-Mott *et al.* (31) (Fig. S1C, movie S3, S4). Using the definition of hydraulic resistance (eq. 2), we designed a range of different geometries (Fig. 1C) that allowed us to explore the putative role of barotaxis in Dd. The bifurcating branches harbour asymmetric features 10 μm downstream of the bifurcation so that no geometrical factors would influence cellular curvature before the decision-making process occurred. In Figure 1D we show how the directional bias (*i.e.* the percentage of cells migrating towards the arm of least hydraulic resistance), changes as a function of the ratio in hydraulic resistance between the two bifurcating arms, for each of the designs shown in Figure 1C.

Analysis of the movement of cells into these channels shows that there is a significant tendency of the cells to move in the direction of lower hydraulic resistance and that this tendency increases with an increase in difference in hydraulic resistance (Fig. 1D), implying that Dictyostelium cells are able to respond to small changes in hydraulic resistance. A closer examination of the results however shows that there is no simple relationship between hydraulic resistance and choice of direction. No statistically significant difference is present between the geometries 2 and 6, where R_h_ = 2 and 8, respectively. On the other hand, a significant difference in the directional bias among the three different geometries (4, 5 and 6) displaying the same hydraulic resistance ratio (R_h_ = 8) was found.

We did not find any evidence for a ‘directional memory’ (30) in our experiments that could complicate the interpretation (Fig. S2). Rather, each cell explores both bifurcating paths at every consecutive bifurcation (movie S1,3,4).

We cannot rule out that small differences in chemotactic signals between the two arms also play a role in these results. In our chip design, the higher concentrations would be expected in the channels with a shorter or wider pathway and thus align with lower hydraulic resistance. Although we assume that there exist at most only small differences in chemo-attractant concentration between the two channels coming from the bifurcation we cannot rule out the residual differences in chemo-attractant concentration that exist at the time of decision making. Therefore, it is formally impossible to conclude, based on experiments using these geometries, whether the cells preferentially respond to a residual chemo-attractant concentration or a difference in hydraulic resistance or possibly a combination of both.

### Decoupling chemical signal from hydraulic resistance

Since the above results could not totally discriminate between the effects of chemotaxis and barotaxis we decided to implement a new design to investigate the competitive effects of chemical cues and hydraulic resistance. We introduced the microchannel topology shown in figure 2A. In this geometry, one of the bifurcating channels is connected to the chemo-attractant source, while the other arm with a low hydraulic resistance feeds back into the “cell loading channel” that does not contain chemo-attractant. This geometry allows to decouple the chemotactic and barotactic stimuli (i.e. “up” the chemical gradient or “down” the hydraulic resistance gradient). The resulting response was unambiguous, 100% (77 out of 77) of cells, analysed over seven independent experiments, migrated up the chemical gradient, despite the hundred times higher resistance of the upper narrow path (movie S5). This held true when the chemotactic gradient of cAMPS was varied 50-fold (20 nM, 100 nM and 1μM). This observation clearly demonstrates that chemotaxis towards cAMP gradients overrides barotaxis in Dictyostelium discoideum in these conditions.

**Figure 2.**
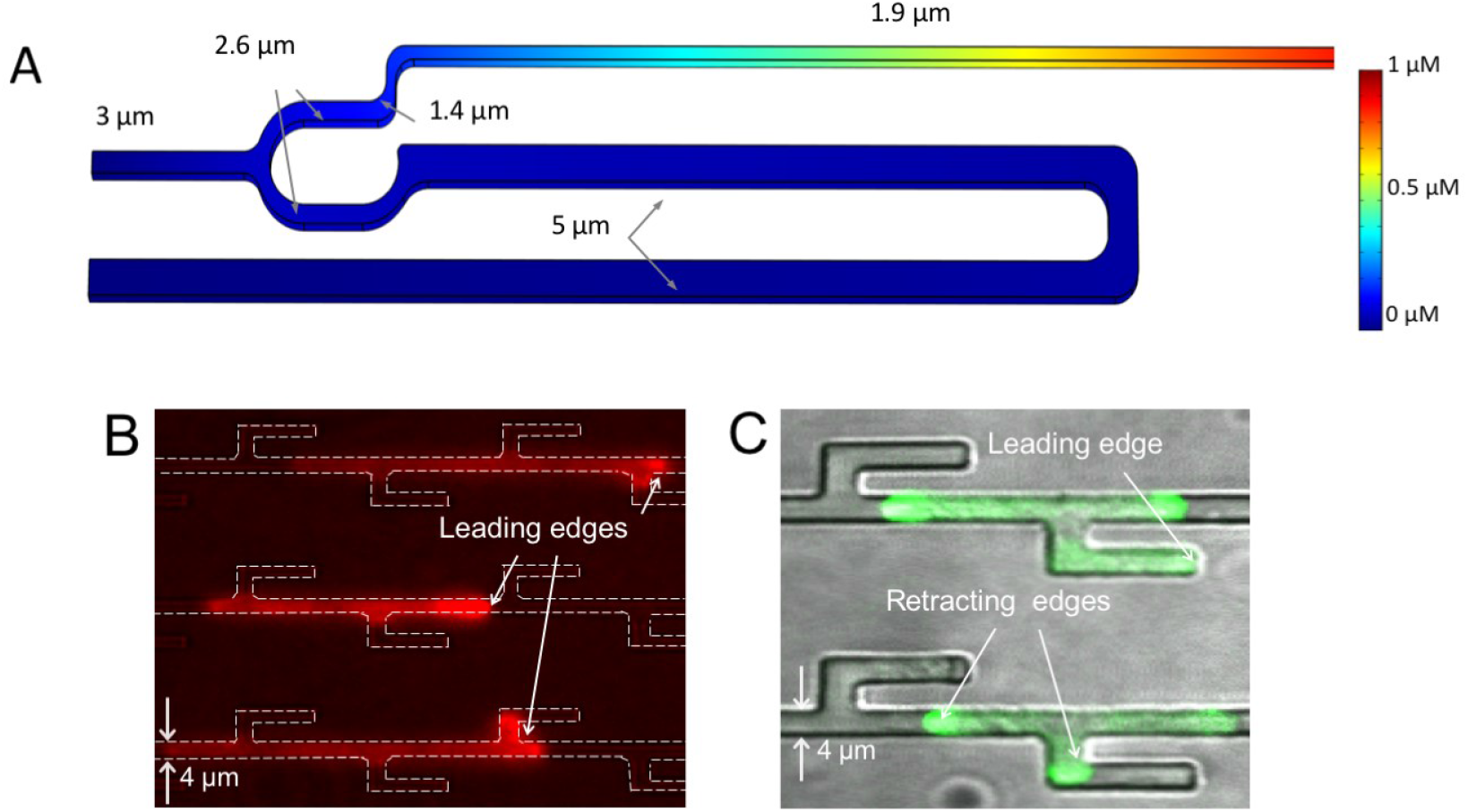
Decoupling barotactic and chemotactic response. (A) Schematic of the migration channel topology that allows for decoupling the chemical signal from the hydraulic resistance. Only the narrower arm joins the cAMPS loading channel (on the right-hand side, not visible), whereas the wider and longer arm goes back into the loading channel where cells are seeded. Cells can therefore either migrate up the chemical gradient or towards the wider channel, where the hydraulic resistance is hundred times lower. The cAMPS distribution across the system is calculated using Comsol. We assume pure diffusion with a concentration of 1 μM at the right side of the channel and 0 μM at the left had side, which in the chip is connected to the cell loading channel. (B, C) Examples of cells actively protruding into dead-end channels. Actin filaments are labelled with RFP-LifeAct in (B) and Myosin II with GFP in (C).

To further test whether *Dictyostelium discoideum* cells respond to hydraulic resistance we designed a migration channel where one of the arms of the asymmetric bifurcations is a dead-end microchannel, and hence in theory has an infinite hydraulic resistance. Many cells exhibited the ability to partially or fully penetrate these dead-end branches (Fig. 2B, C, Fig S3, movie S6). In the case of total or partial penetration, the cells appear smaller than the cross section of the channel and therefore the fluid in front of them can be displaced around the cell through the imperfect seal between the cell and the channel wall. In the cases, where cells do not invade blind channels, the leading edges seem to plug the channel completely and cannot invade the dead-end branches (Fig. S3). To further investigate this aspect, we added a fluorescent dye FITC Dextran (70kD) into the loading channel at the right-hand side of the chip, together with the chemoattractant. We observed that as the cell protrudes into the blind channels the fluid was displaced around the leading edge, while there was no evidence for micropinocytosis as has been reported for immature dendritic cells recently (32). These results appear to indicate that that cells are only able to migrate into dead-end channels under conditions where they can deform sufficiently to shunt some fluid between the wall and membrane to generate space to move into. But even under these likely high hydraulic pressures, the cells will still explore these channels. These observations also do not support the idea that barotaxis is a key driver in the decision to extend a pseudopod or not and argue against barotaxis as a dominant mechanism governing Dd cells' directional choice.

### Splitting vs Non-Splitting

When a cell encounters a bifurcation, it often extends a pseudopod in both channels. Labelling of the actin cytoskeleton with Lifeact shows that these leading pseudopods are, as expected, enriched in filamentous actin (Fig. 2B), while Myosin II-GFP strongly localises at the uropod and at retracting ends (Fig. 2C, movie S1–S3). Interestingly, we observe that some cells did not split in two pseudopodia at the bifurcation shown in Figure 2A, but instead were able to directly move into the channel connected to the cAMPS source. The splitting occurrence increased with the cAMPS concentration, at the highest cAMPS concentration tested 80% of the cells split (Fig. 3A). This poses two questions: 1) why do the cells extend a pseudopod in the wrong direction and 2) when and how do they make the decision to retract?

**Figure 3.**
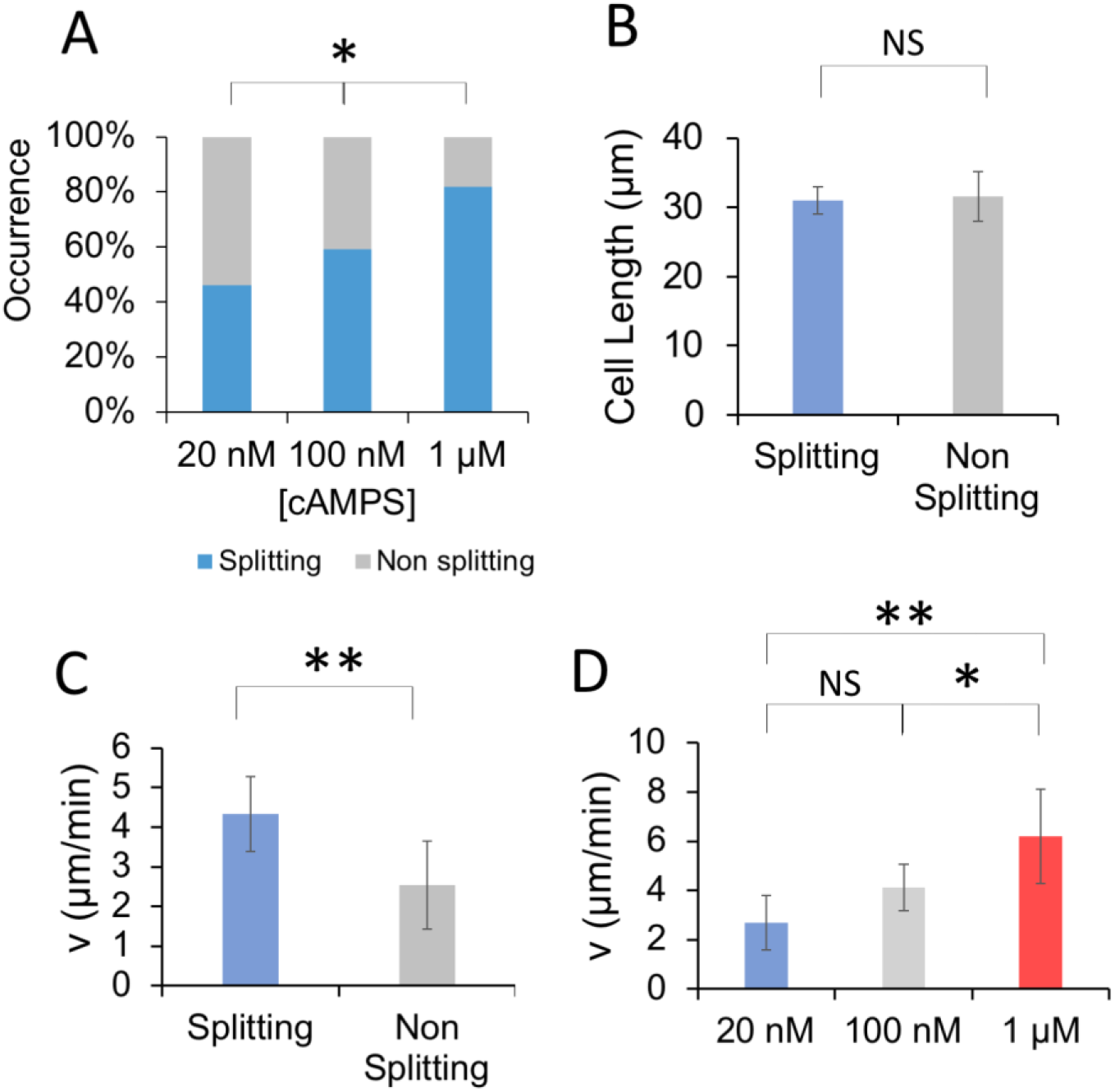
Relationship between cell properties and splitting behaviour at the bifurcation at the asymmetric bifurcation in Figure 1C. (A) Relationship between the cAMPS concentration and the cell splitting. (p=0.039, using 2×3 Fisher Exact Test). (B) Relationship between cell length and splitting behaviour. Median values are shown. p=0.008 (C) Relationship between splitting behaviour and the velocity of the cells in the straight channel upstream of the bifurcation. Median values are shown. (D) Relationship between the cell speed and the cAMPS concentration for the splitting subgroup. NS > 0.06, *=0.06, **=0.02. Kruskal-Wallis Test was performed in B-D. Error bars represent 95% confidence interval.

We sought to better understand what parameters affected these behaviours. We found that splitting did not depend on cell size (Fig. 3B), but we did notice a significant correlation between the speed of cell before entering the channel and the frequency of splitting (Fig. 3C). We also determined that cell speed increased with the cAMPS concentration in these conditions (Fig. 3D) similar to previous reports that showed that cells under non confined conditions showed an cAMP dependence on migration speed (33). Thus, the recorded dependence of migration speed on cAMPS concentration likely underlies the observed dependence of splitting on cAMPS concentration (Fig. 3A). It is well established that the formation of new pseudopods during normal and chemotactic motion requires localised actin polymerisation at the leading edge and that this contributes to the driving force necessary for extension and migration. We observed that cells migrating in long straight channels tended to exhibit a persistent localisation of polymerised actin at the leading edge and to a lesser extent at the trailing edge (Fig. 4A). Within a given cell population there was a considerable individual variation in the amount of polymerised actin at the front, which were expected to reflect the observed variations in migration speed. Measurement of the linear extent of the actin polymerisation zone at the leading edge for each cell correlated well with the speed of the cells (Fig. 4B). Since faster moving cells also split more readily, this suggests that cell specific differences in internal actin dynamics may determine whether splitting occurs (Fig. 4D-F).

**Figure 4.**
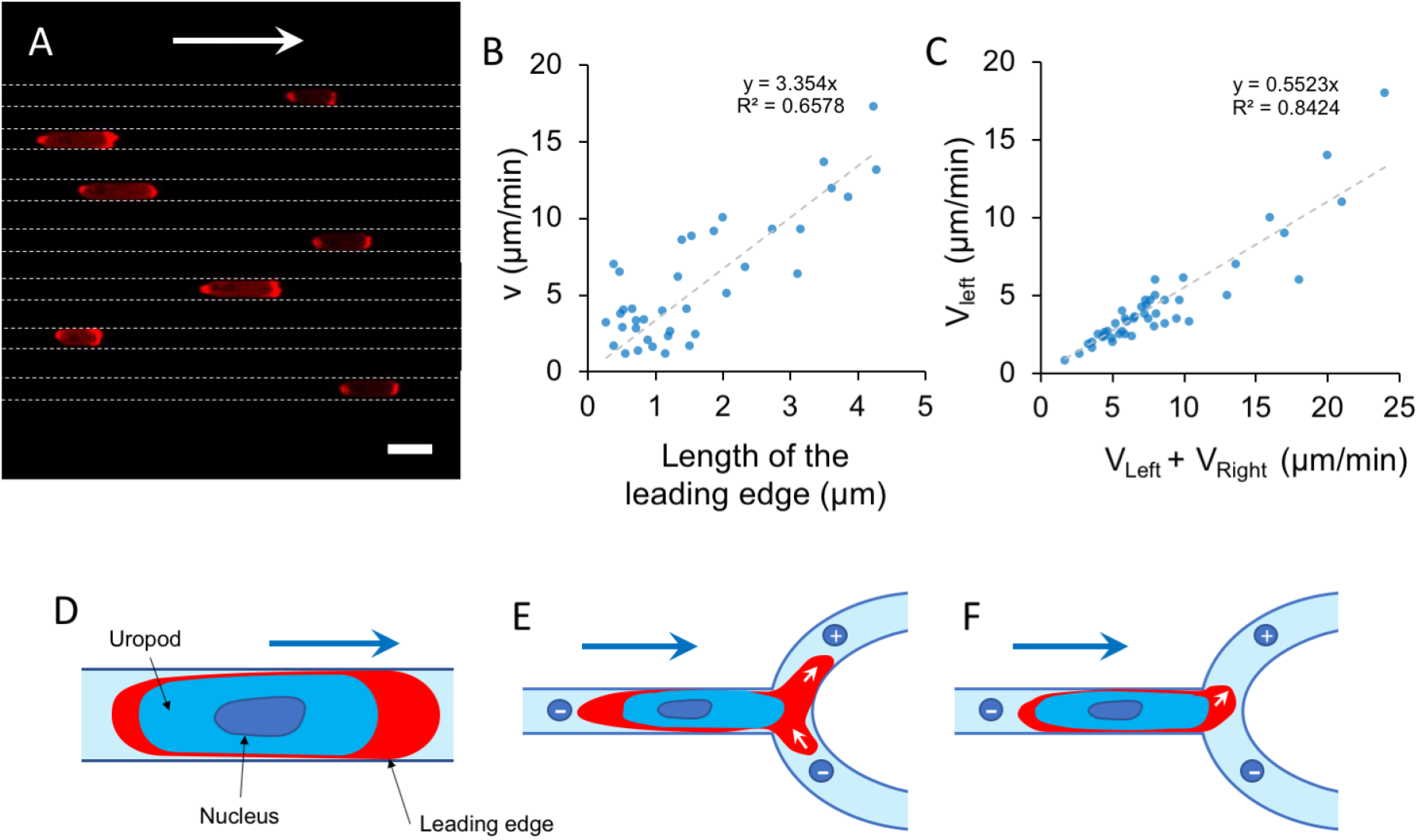
Relationship between the size of the actin rich leading edge and the splitting behaviour at the bifurcation at the asymmetric bifurcation in Figure 2A. (A) Example of cells migrating in straight channels with characterise by a cross section of 5 × 2 μm^2^. Actin filaments, labelled with RFP-LifeAct, primarily localises at the leading edge and, in lower amount, at the uropod. Scale bar is 10 μm, (B) Relationship between the length of the actin rich leading edge and the velocity of 37 cells migrating in straight channels with cross section 3 × 2 μm^2^. (C) Relationship between the speed of the leading edge along the path of positive cAMP gradient (V_left_) and the sum of the average speeds of the pseudopodia protruding in both bifurcating channels (V_left_+V_right_). Data are pooled from 2-4 experiments at different cAMP gradients (20 nM, 100nM and 1μM). See fig S4 for data at the individual concentrations. (D) Schematic representation of the actin distribution of migrating cells confined in microchannels. (E) Schematic representation a fast moving cell with large leading edge, which tends to split. (F) Schematic representation of a slow moving cell with small leading edge, which tends to migrate straight towards the source of chemoattractant.

Next we aimed to better understand when and how cells make the decision to retract the pseudopods along the developing negative cAMPS gradient. We had noticed that not all cells bifurcate pseudopods and that among the ones who did there was a considerable variation in the length of extension before the decision to retract was taken. To characterise this behaviour further we have performed a quantitative analysis of the dynamics of bifurcation. We have measured the length and average speed of extension of the pseudopods protruding inside the bifurcating arms up to the point where the pseudopod moving down the cAMPS gradient stopped moving and started to retract. We found that there exists a very strong positive linear correlation between the average speed of extension of the pseudopod up the gradient and the sum of the average speeds of extensions up and down the cAMPS gradient. The slope of this dependency is ~0.55. Interestingly the slope does not depend significantly on the steepness of the cAMPS gradient across the cell, as shown in Figure S4. Since during cell migration there is no significant inertia, average pseudopod expansion speed is proportional to the net local driving force. These data therefore strongly suggest that a pseudopod is retracted when a critical imbalance in the driving forces in the competing pseudopods is reached.

## Discussion

We have tested whether Dd cells would respond to asymmetric hydraulic resistance conditions by confronting them with a variety of microfluidic channels of different topologies as has been reported for neutrophils (31). There was definitively a tendency for the cells to preferentially migrate into channels of lower hydraulic resistance. However, the interpretation of these results was complicated by the fact that we could not completely decouple the chemotactic stimulus from the barotactic stimulus. Lower hydraulic resistance coincided with a shorter or wider path to the cAMP reservoir, which could potentially result in slightly higher cAMP concentrations, at least transiently. As a result of this we cannot completely rule out that cells did not respond by chemotaxis rather than barotaxis.

The system developed by Prentice-Mott *et al.* (31) is also not capable of decoupling the contribution of the hydraulic resistance and the chemical gradient as in their device the bifurcating arms of least hydraulic resistance would also correspond to the path with steepest chemical gradient. Although Prentice-Mott *et al.* found that the directional bias was the same both in the presence and in the absence of gradients of chemo-attractant (fMLP), a possible caveat could still be that these cells undergo *autologous chemotaxis,* as previously observed in cancer cells and neutrophils (34, 35).

The mechanism of hydraulic pressure sensing is also rather mysterious. It can be estimated that, to be barotactic, Dd cells should be able to measure very small differences in pressure at their leading edges to respond to the local differences in hydraulic resistance. In fact, the force necessary to push a column of water through a microchannel is given by the capillary force (36):

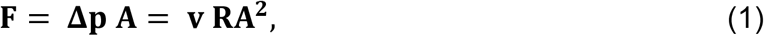

Where *v* is the cell velocity, *R* is the hydraulic resistance of the channel and *A,* the cross-section. Using the definition of *R* given in the Eq. 2 and the following values: *L* = 250 μm, *w* = 3 μm, *v* = 8 μm/min, *μ* = 1 mPa • s, *h* = 2.5 μm the estimated force is in the order of 0.5 pN. Therefore, the force difference between the two arms of a bifurcation is in the order of a few hundred femto-Newtons. These forces are very small, even smaller than that generated by a single actin filament that polymerises at the leading edge (in the range of a few pico-Newtons (37, 38).

To better understand the directional bias exhibited by cells moving through asymmetric bifurcations, we developed a microchannel possessing an asymmetric bifurcation aiming to decouple the two main environmental stimuli competing in the system: the chemical gradient and the hydraulic resistance. This topology allowed us to show that all cells analysed migrated towards the positive chemical gradient, rather than moving into the arm of hundred times lower hydraulic resistance. Our results show that chemotaxis overrides barotaxis in Dictyostelium, if barotaxis does exist in Dictyostelium.

These finding that Dd cells do not respond strongly to hydraulic resistance is also supported by the observation that many cells are able to move into dead-end channels, despite the theoretical infinite hydraulic resistance of these channels. To explain how cells can move into dead-end channels a number of different mechanisms have been proposed. It has recently been demonstrated that tumour cells confined in microchannels display a polarised distribution of ion pumps (Na^+^/H^+^) and aquaporins in the cell membrane. This is assumed to create a net influx of water at the leading edge and a net outflow at the uropod driving the cells motion and represents an alternative to the actin-driven motion (*‘Osmotic Engine Model*’ (39)). Recent work by Moreau et al. (32) suggested that immature dendritic cells (DCs) cells could cope with high hydraulic resistance via a high level of macro-pinocytosis, that allows fluid transport across the cell and thereby exploration of dead-end channels (40). Mature DCs down-regulate macropinocytosis and lose the ability to penetrate dead-end channels. Although growing Dd cells show a high level of macro-pinocytosis, no uptake of labelled dextran from the fluid via macropinocytosis was observed during our experiments with starving Dd cells, making the latter mechanism an unlikely explanation (Fig. S2). The ability of Dd cells to move into the dead-end channels is likely due to the fact that the fluid confined inside the blind channels in front of the protruding leading edge of the cell manages to leak between the cell and the channel wall, as the cell protrusion advances into the dead-end branch (Fig. S2). Taken together, we consider it is unlikely that starving Dd cells show a significant barotactic response but do as yet not rule it out completely.

Directed and conditional pseudopod extension and retraction is key to chemotactic mechanism. The *compass-based* model predicts that cell first senses the direction of the attractant gradient, polarises followed by the extension of a new pseudopods in the direction of the gradient (41). Movement in steep gradients can be explained as the persistent extension of pseudopods in the correct direction, largely alleviating the need for pseudopod retraction. However, previous work by Insall and co-workers (42, 43) has shown that Dd cells migrating in shallow gradients of cAMP move via a mechanism that involves the generation of bifurcating pseudopods at the leading edge of the cell. The pseudopods appear to be generated in random directions, but the pseudopod pointing towards higher concentration of chemoattractant is stabilised, while the pseudopods pointing in other directions is retracted. This results in a slow but reproducible turning of the cell in the direction of the gradient. Although this decision to extend or retract is clearly dependent on the gradient of cAMP, it remains unresolved at which stage and by what mechanism the cell decides to retract a pseudopod pointing in a less favourable direction.

Our experiments have shed some light on the decision-making process on pseudopod extension and retraction. We found that fast moving cells split in two when reaching a bifurcation and the pseudopod protruded inside the arm of low cAMP extended for a short time before stopping and retracting, a process that involved the recruitment of myosin II. Slower moving cells did not split and moved straight into the correct channel. We also found a clear relationship between the extent of the zone of filamentous actin at the leading edge and the speed of movement, faster cells having a larger actin filamentous acing rich zone. We also confirmed that under our conditions of high confinement cells experiencing higher cAMP gradients migrate faster than cells experience smaller cAMP gradients. Therefore, it seems likely that high levels of cAMP-dependent actin polymerisation, results in a more active and larger actin polymerisation zone at the leading edge, which may increase the probability of splitting at a bifurcation.

The measurement of pseudopod retraction dynamics may shed some light on the underlying mechanism. We demonstrated that the average velocity of the pseudopodia protruding up the cAMPS gradient is linearly proportional to the sum of the average velocities of the competing pseudopodia protruding up and down the cAMP gradient. The pseudopod pointing down the gradient (along the path with 100 times lower hydraulic resistance) retracts when the average velocity of the pseudopod pointing up the gradient is 20% larger than its own average velocity. This threshold is perhaps surprisingly independent of the magnitude of the cAMPS gradient over the cell. Since speed in this overdamped system is proportional to the local driving force, this implies that the decision to retract is independent of the absolute difference in driving forces between the two ends, but only depends on the steepness of the gradient. It has been argued that during neutrophil migration a cortical tension gradient along the cell might be responsible for maintenance of cell polarisation, i.e. tension prevents initiation of new pseudopods, but it has been difficult to estimate how large this tension gradients would be (44, 45).

Our results suggest that this tension gradient has to be 20% or larger be able to suppress a secondary leading pseudopod. The mechanism underlying tension dependence will have to be addressed in future experiments.

## Material and Methods

### Microfluidic Device

The structure of the microfluidic device is similar to that used in other studies on neuronal cultures (46) and tumour cell invasiveness and migration (28), and is characterised by the presence of two main loading channels, each of which connects two cylindrical reservoirs that act as infinite source and as sink, respectively. The two ‘loading channels’ are bridged by ‘migration channels’ having a cross-sectional area that is much smaller than that of the loading channels. The 3D schematics of the device (Fig. 1A). The cell suspension is loaded in one side of the chip, whereas the other symmetric half is loaded with the chemoattractant (cAMPS).

The hydraulic resistance of a duct can be defined as *R_h_= ΔP/Q*, where *ΔP* is the pressure drop and *Q* the flow rate (47). This formula can be considered the hydrodynamic analogous to Ohm’s law *R= ΔV/I*, where *ΔV* is the voltage and *I* the current through a wire. In the case of micro-channel with a rectangular cross section the hydraulic resistance can be expressed as:

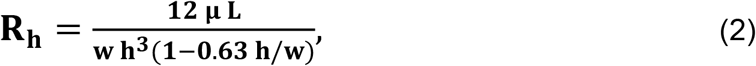

Where *w* is the width of the channel, *L* is its length, *h* is its height and *μ* is the fluid dynamic viscosity (47). The migration channels in Figure 1C have been designed using Eq. 2 by varying the length and cross-sectional diameter of the channels. In all cases the height of the channels was fixed at 2.5 μm.

### Device fabrication

The design of the mould is carried out in QCAD professional (version 3.9.8.0). The moulds of the microfluidic chip were developed using standard photolithographic techniques (48). First, a 2.5 μm high film of the photopolymer SU-8 2002 was spread onto a six-inch silicon wafer (IDB Technologies) via spin coating to produce the first layer of the mould. A first high-resolution photomask was used to transfer the features of the migration channels through UV illumination. A second 80 μm high layer of SU-8 2075 was then spun on the wafer and the features of the bigger loading channels were aligned with those of the first layer through alignment marks and transferred to the second layer via UV illumination. All the areas that were not exposed to the UV light were then etched using a SU-8 developer. Replicas of the patterned mould were obtained by soft lithography using polydimethylsiloxane (PDMS) (Sylgard 184 Silicone Elastomer Kit; Dow Corning Corporation). The PDMS base and curing agent were mixed at 10:1 ratio, degassed in a vacuum desiccator, and poured on the mould. After baking for 6 hours at 80° C the solid PDMS was separated from the mould and ports were punched through the inlets and outlets regions using 8 mm biopsy punchers (Harris Uni-Core). Patterned PDMS was reversibly bonded to standard microscope glass slides (VWR) exploiting the Van der Waals forces that generate at the interface between the PDMS and the glass.

### Cell culture

Dictyostelium discoideum cells (strain A×2) were used. Cells were grown in shaken suspension in HL5 medium using glass flasks at 22° C. Cells were harvested when the cultures reached ~4×10^6^ cells/ml cells by centrifugation (400 g for 2 minutes). Cells were washed twice and re-suspended in KK2 buffer at the concentration of 1×10^7^ cells/ml and starved for 4.5 hours while subjected to periodic stimulation with 10^−7^ M cAMP (49) [46]. To visualise actin dynamics a strain transformed with Lifeact RFP was used, to visualise myosin dynamics a myosin II heavy chain GFP knockin strain was used (Singer and Weijer, unpublished).

### Cell loading

After starvation, cells were washed twice and suspended in KK2 at a final concentration of 1×10^7^ cells/mL. An aliquot of 200 μL cell suspension was loaded inside one of the two inlets of the microfluidic device using a pipette. The opposite inlet was filled with 200 μL of non-hydrolysable cyclic AMP analogue cAMPS at increasing concentration: 20 nM, 100 nM, 1 μM. After loading the system with the cell and chemoattractant solutions, it becomes subjected to a transient flow directed from the source to the sink reservoirs. The flow lasts until the hydrostatic pressure at these two ends of the loading channels is balanced. At the equilibrium, a linear spatial gradient of chemoattractant establishes inside the bridging channels (Fig. S1), mainly by diffusion, as previously shown by Abhyankar and co-workers (50). The symmetry guaranteed by the microfluidic device allows for a very good equilibration of the system. Cells adhere to the substrate and those that sit in the proximity of the entrance of the migration micro-channels readily sense the cAMPS and move up the gradient, while undergo a drastic shape modification to adapt to the mechanical confinement. The choice of using the non-hydrolysable cyclic AMP analogue cAMPS was based on the fact that upon exposure to the cAMP gradient each Dd cells secrete intracellular cAMP at a concentration comparable to that imposed by the system (1 μM) (Tomchik et al. 1981), that would have affected our analysis. Moreover, Dictyostelium cells express and secrete phosphodiesterase (PDE), which degrades cAMP into 5’-adenosine monophosphate (51). This means that the local concentration around the single cell would have been the result of these two competing effects and cells that migrate close to each other could interact. The use of cAMPS simplifies the chemical landscape cells are exposes to. At the end of each experiment the glass cover slip was separated from the PDMS chip and both surfaces cleaned with ethanol 70% and stored for following experiments. Additionally, to maintain the chemical gradient steady, the four reservoirs were covered using small pieces of PDMS.

### Imaging, image analysis and statistical analysis

The cell motion was recorded at room temperature on a Leica SP8 confocal microscope equipped with a 10×, 0.3 N.A. plan apochromat objective lens. The length of the splitting pseudopodia was quantified manually using ImageJ by measuring the distance of the extent of each pseudopod in both bifurcating arms, relative to the centre of the bifurcation. Similarly, the length of each cell and of the actin polymerisation zone was estimated manually using ImageJ. The speed of the cells and pseudopodia was measured using ImageJ, via kymograph analysis. The inclusion criteria for our analysis were that cells had to travel directionally towards the higher end of the chemical gradient. In the case multiple consecutive cells were migrating inside a channel, only the first cell was considered. To characterise how cells interacted with the surrounding fluid, FITC-dextran (FD-70, Sigma-Aldrich) was loaded into the right-hand side reservoirs along with cAMPS. Data analysis was conducted in Excel for Mac 2011, Matlab R2018b and Python.

### Numerical simulation

To estimate the local concentrations of cAMPS cells were exposed to at the bifurcating microchannels we simulate the transport of cAMPS through the migration channels using the software Comsol Multiphysics 5.2b (Comsol, Inc.). The transport of cAMPS was assumed to be driven by diffusion. We calculated the equilibrium solution of the 2D diffusion equations for the actual topology of the bifurcating channels using the module *‘Transport of Diluted Species’*. *Constant-concentration* boundary conditions were imposed at the two ends of the channel, the values at these positions were set to be c = 1 μM and c = 0 μM, respectively, as in the actual experiments. The boundary conditions were set as *‘no-flux’* at the edges of the channel. The initial conditions were 0 μM throughout the channel. The value of the cAMP diffusion coefficient, *D*, in water is 4.44×10^−6^ cm^2^/s (52).

## Acknowledgments

We acknowledge Gail Singer for help with the Dictyostelium strain generation and culture, Steven Neale (University of Glasgow), Michael MacDonald, Paul Campbell, Serenella Tolomeo and Philip Murray (University of Dundee) for helpful discussions and constructive comments. The microfabrication of the microfluidic devices was carried out in the cleanroom facility at the Division of Physics of the University of Dundee (UK). This work was supported a PhD studentship of the Engineering and Physical Sciences Research Council (EPSRC) and a BBSRC BB/L00271X/1 grant to CJW.

## Supplementary Information

**Figure S1.**
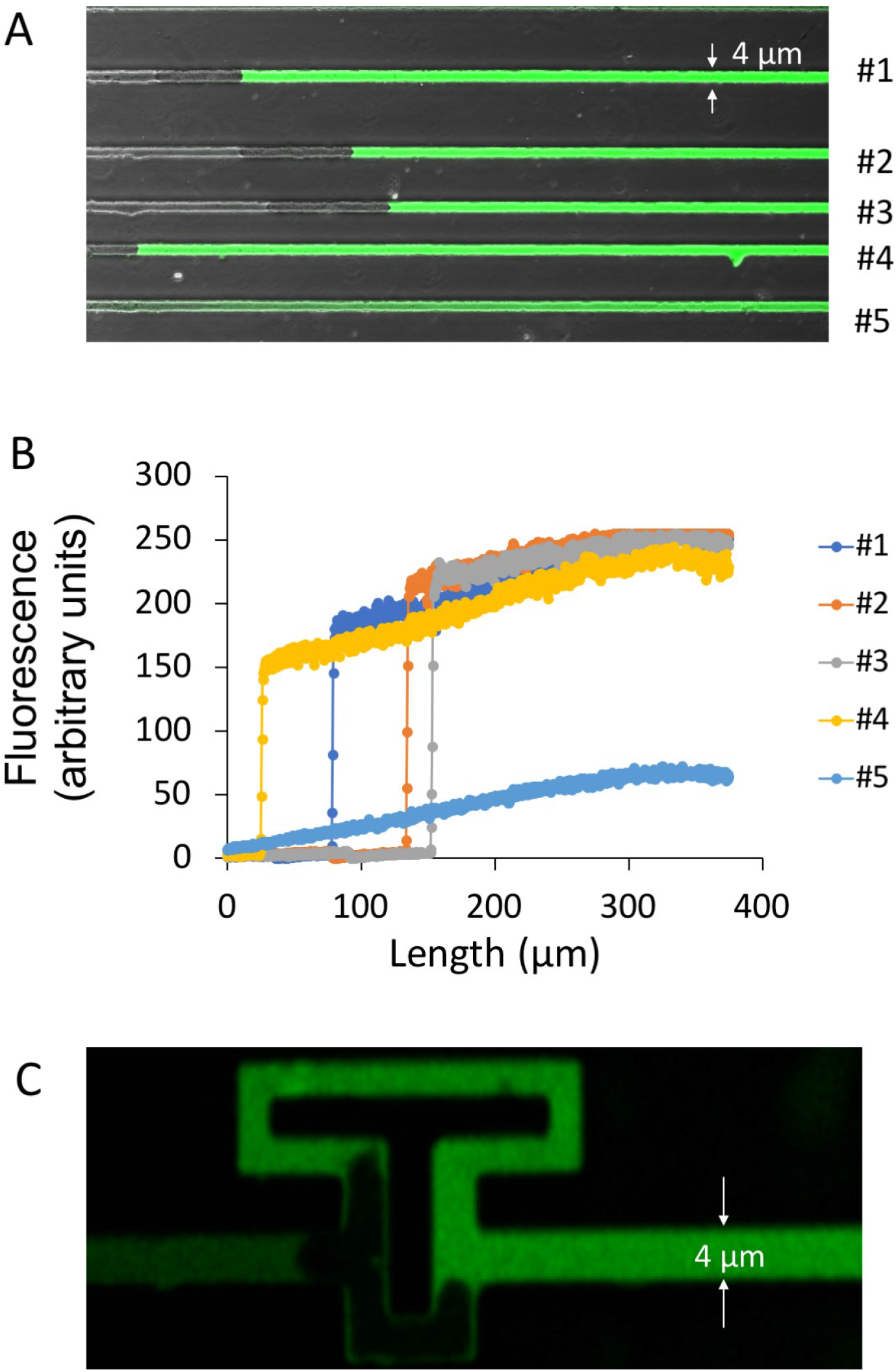
Gradients within channels with migrating cells. A) partial view of migration chip showing the middle 375 μm section of 5 channels. Cells have entered at slightly different times in channels 1-4, while no cell has entered channel 5. The cAMP loading channel was filled with a 100nM cAMPS solution containing 0.1% FITC-dextran to visualise the distribution of a small molecule in this system. B) Concentration profiles of the dextran fluorescence of channels 1-5. Note the steep fluorescence gradient over the length of the cells in channels 1-3 and the graded concentration profile in channel. C)Distribution of FITC-dextran in an asymmetric bifurcating channel in the presence of a cell exploring both arms of the bifurcation.

**Figure S2.**
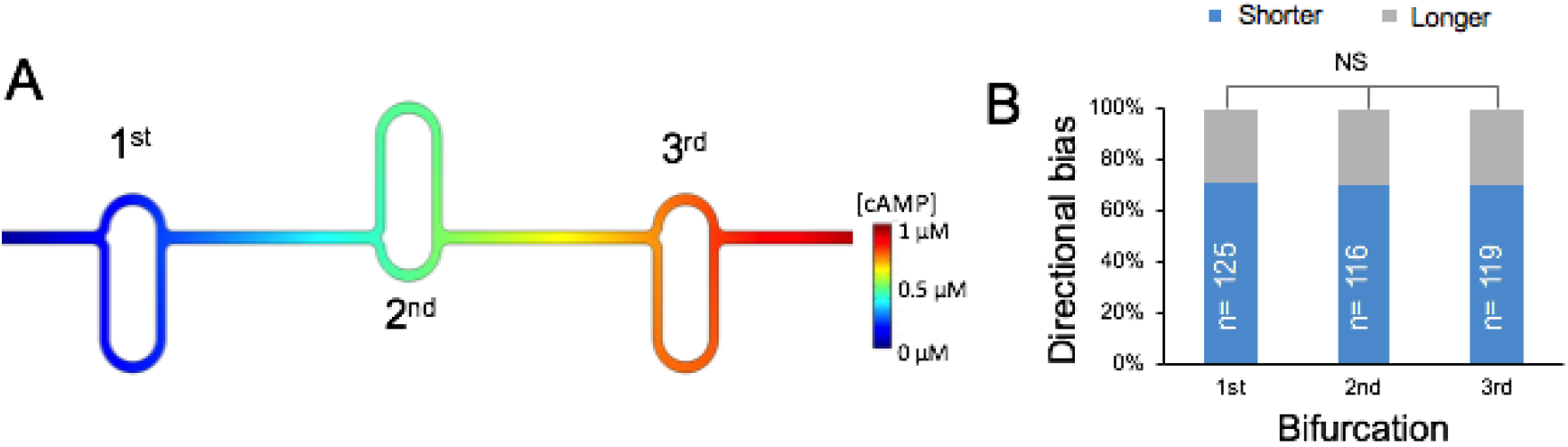
Investigating directional memory. (A) Bifurcating asymmetric channels arranged in series. The position of the longer/shorter arms is inverted at each bifurcation point to account for potential *‘directional memory’.* (B) Directional bias in the case of cells migrating through the channels in (A).

**Figure S3.**
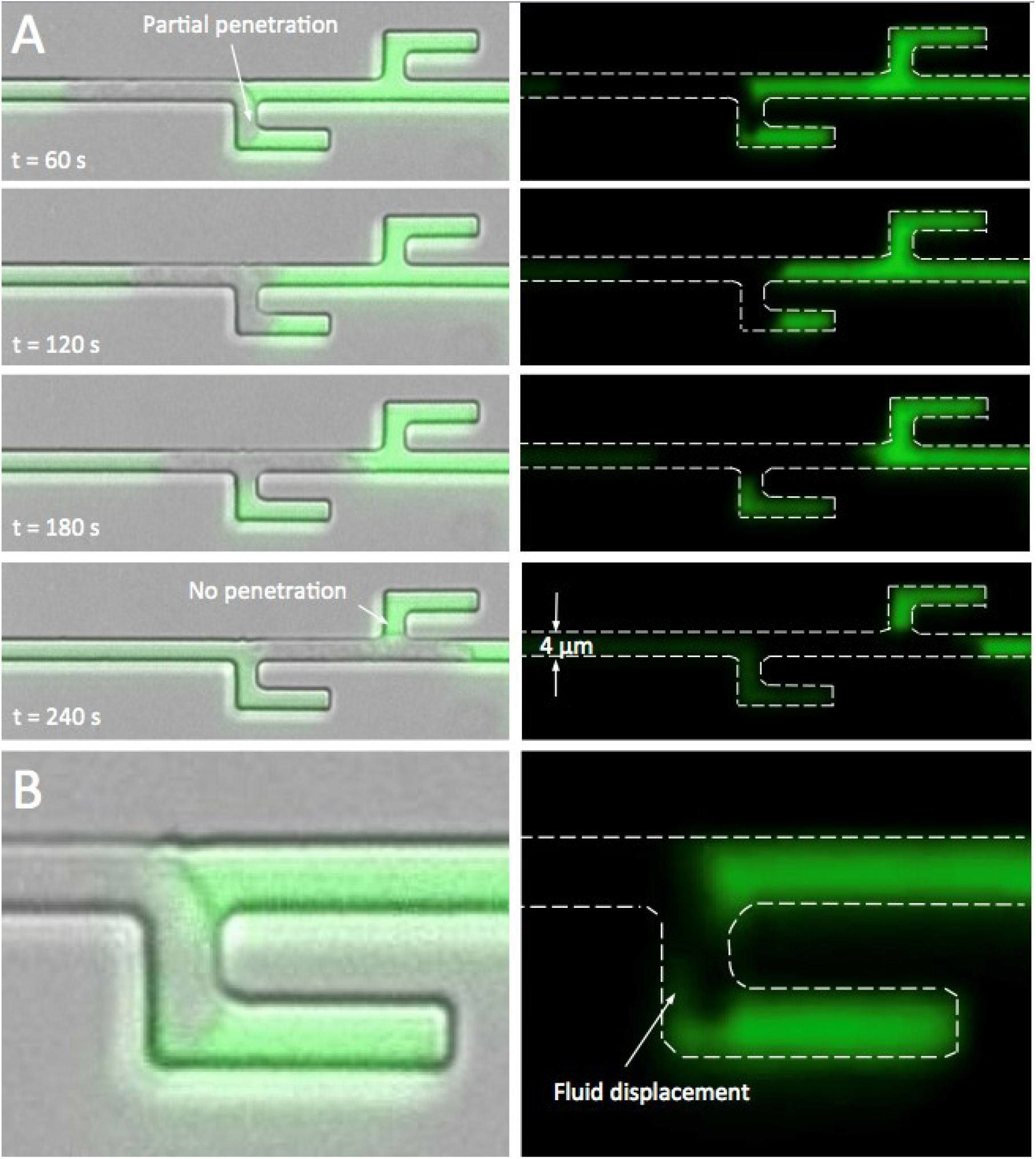
Cell confined in microchannel with alternating dead-end bifurcations. (A) The image sequence shows a Dictyostelium cell that encounters two consecutive dead-ends. The cell protrudes into the first blind arm but not in the second one. FITC-dextran is added to the right-hand side loading channel to investigate whether the fluid is displaced by the cell upon invasion of a dead-end channel. The channel is 2.5 μm high. (B) Close-up of first blind arm shown in the first image in (A). The picture clearly shows a fluid displacement around the advancing leading edge.

**Figure S4.**
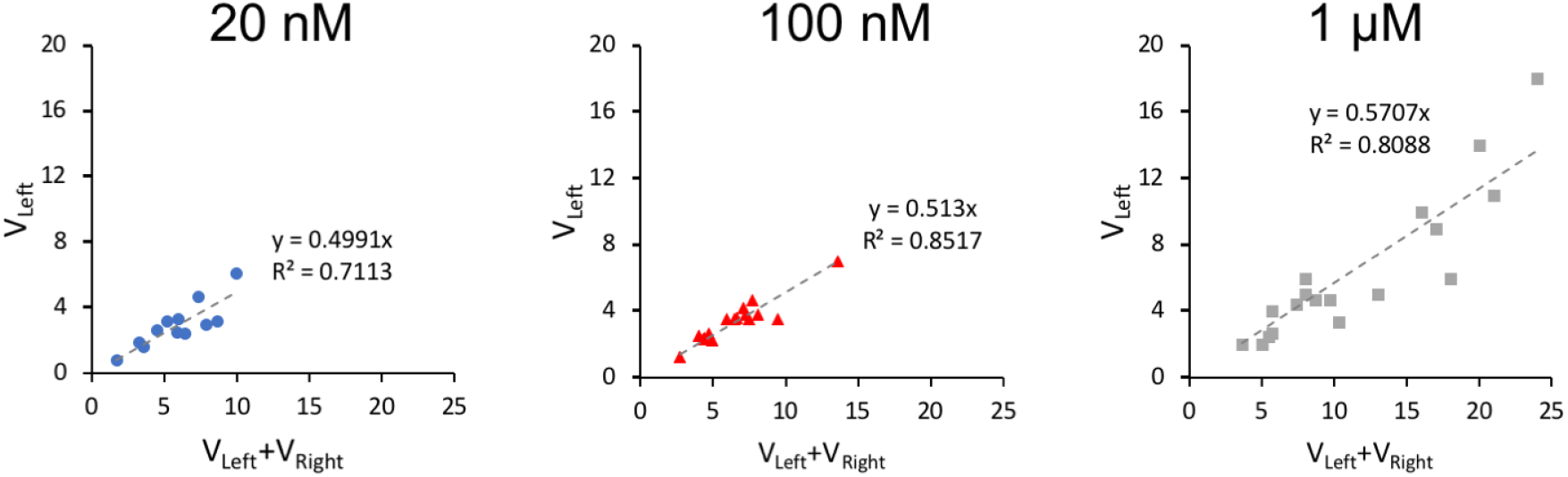
Relationship between the speed of the leading edge along the path of positive gradient and the sum of the average speeds of the pseudopodia protruding in the bifurcating channels shown in Figure 2A.

## Supplementary Movies

**Movie S1.**
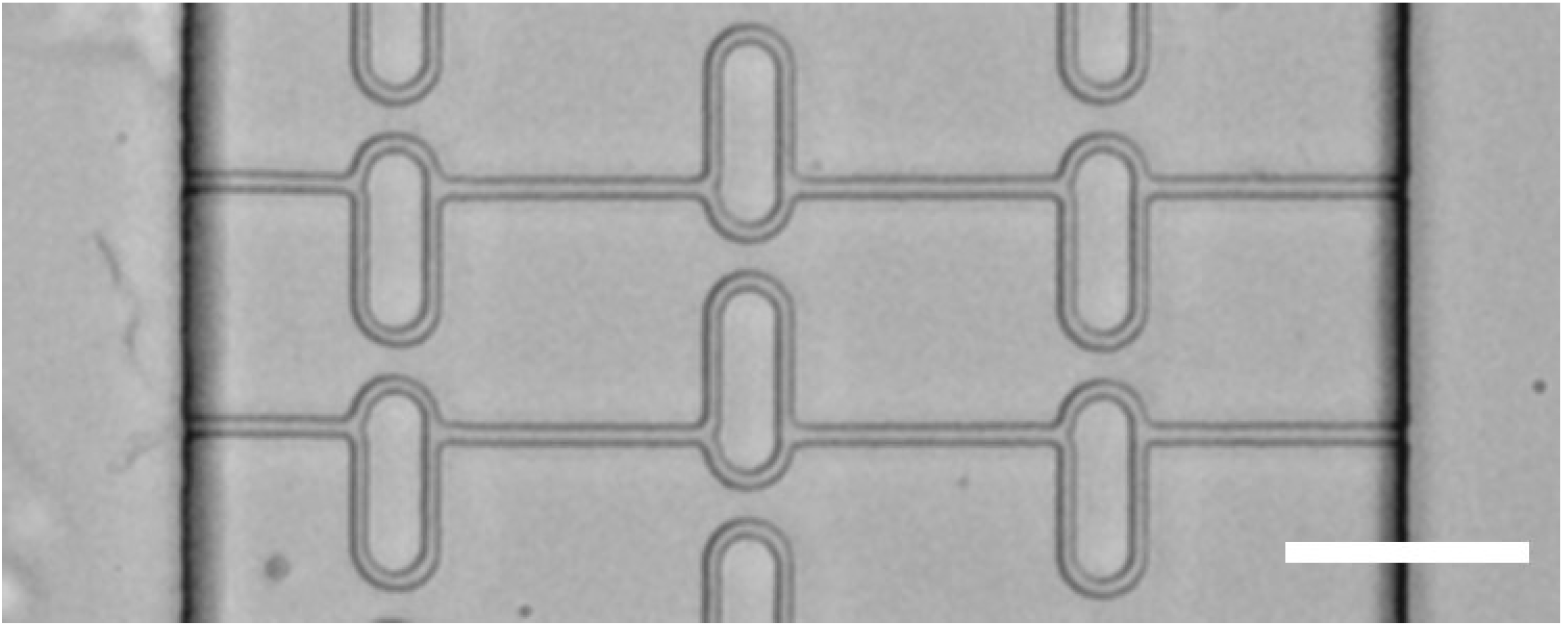
Dictyostelium discoideum cells traverse microfluidic channels in response to a gradient of cAMP the cell loading channel is on the left and the cAMPS loading channels is on the right. The movie is taken at 30 second intervals and played at a frame rate of 10 frames/s. Scale bar is 50 μm.

**Movie S2.**
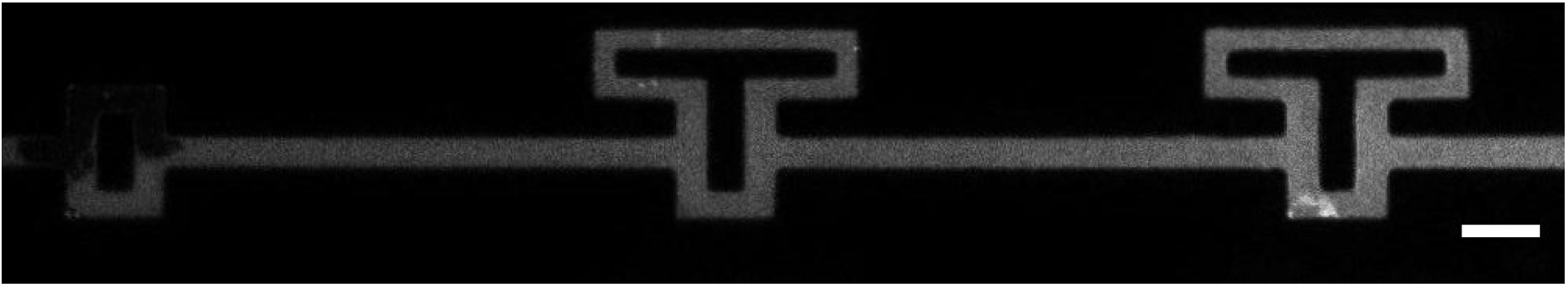
Dictyostelium discoideum cell travels up a cAMPS gradient. The channels cAMP loading channel contains 0.1% fluorescein dextran. It can be seen that there is a steep gradient over the length of the cell. Images were acquired every 30 seconds and the movie is played at 30 frames/s. Scale bar is 10 μm.

**Movie S3.**
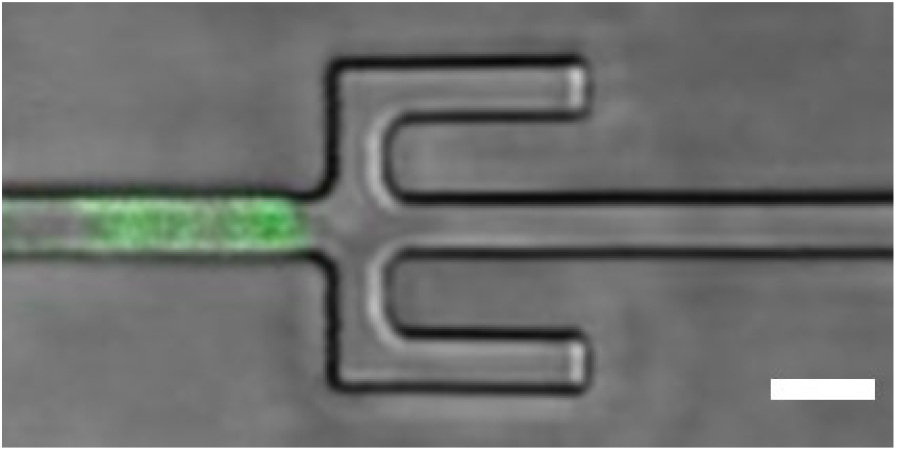
Dictyostelium discoideum cell travelling through a symmetric bifurcating microchannel. The cell generates two almost symmetric pseudopods, one in each arm of the bifurcation and competition between them occurs. The losing pseudopod retracts, and the cell moves towards the exit of the bifurcation. Myosin II - GFP localises at the trailing edge and at the retracting leading edge. The cAMPS concentration increases towards the right-end side. Images were acquired every 30 seconds and the movie is played at 30 frames/s. Scale bar is 10 μm.

**Movie S4.**
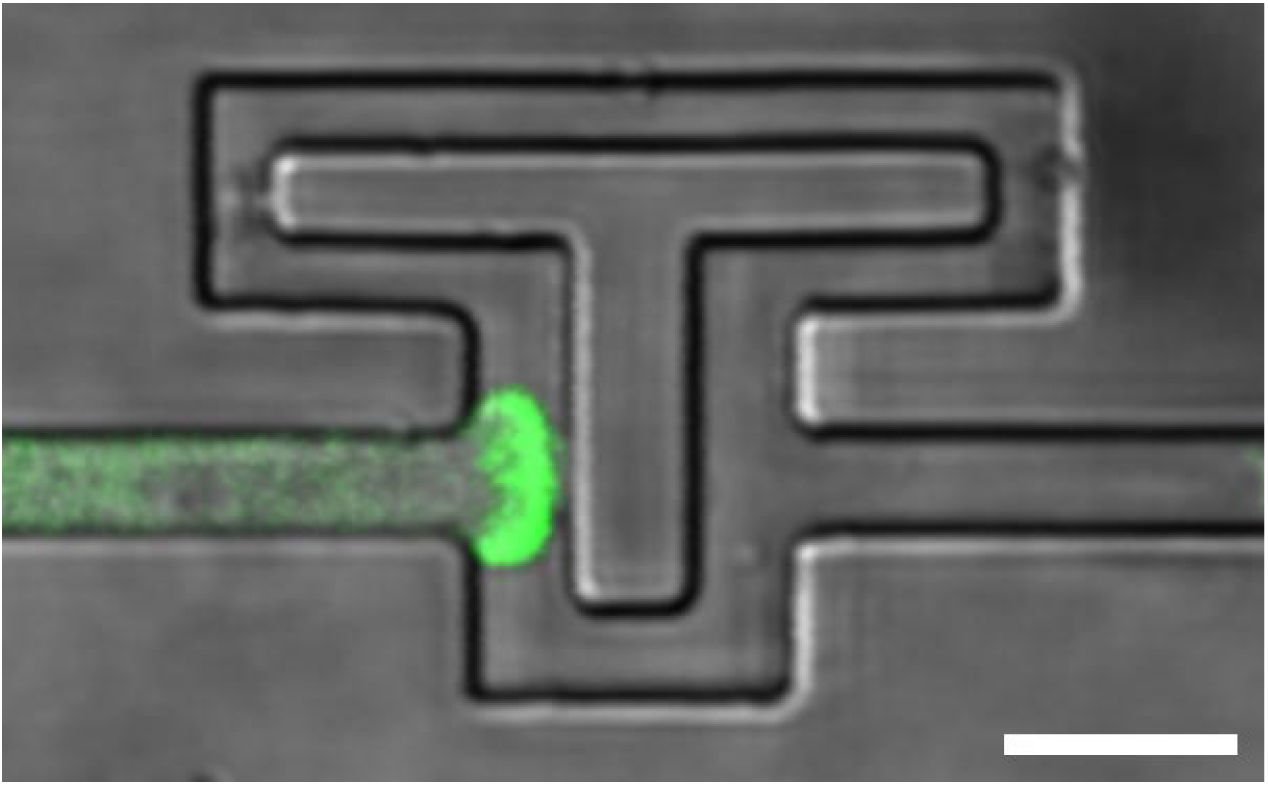
Dictyostelium discoideum cell travelling through an asymmetric bifurcating microchannel. The cell generates two almost symmetric pseudopods, one in each arm of the bifurcation and competition between them occurs. The’winning’ pseudopod splits again where the two paths re-join with the main channel. The secondary pseudopod retracts, and the cell moves towards the exit of the bifurcation. Myosin II - GFP localises at the trailing edge and at the retracting leading edge. The cAMPS concentration increases towards the right-end side. Images were acquired every 30 seconds and the movie is played at 30 frames/s. Scale bar is 10 μm.

**Movie S5.**
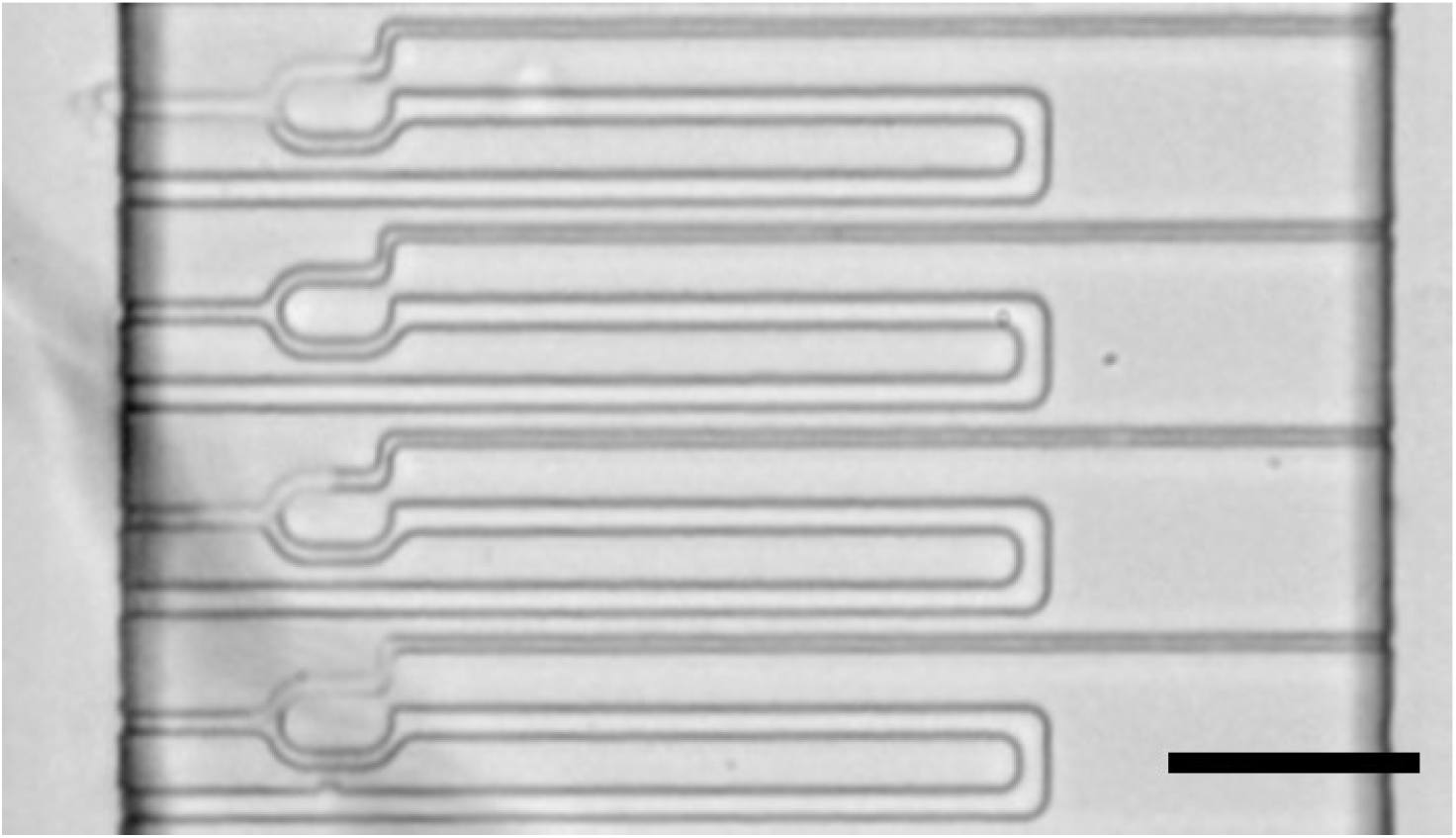
Chemotaxis vs Barotaxis. This channel’s geometry allows us to decouple the chemical signal from the hydraulic resistance. In fact, only the upper arm relates to the source of the chemical signal (cAMPS), while the lower arm has a hundred times lower hydraulic resistance. Images were acquired every 30 seconds and the movie is played at 10 frames/s. Scale bar is 50 μm.

**Movie S6.**
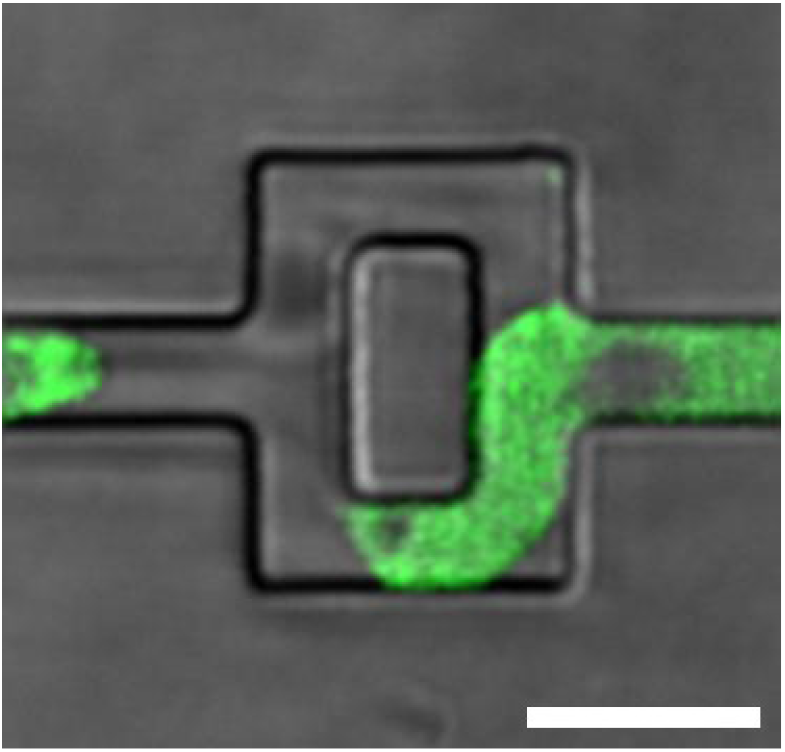
Dictyostelium discoideum cell travels up the cAMPS gradient and navigates through a microchannel with two symmetric bifurcating dead-ends. The cell explores all the possible directions before to move towards the exit of the main channel. Myosin II - GFP localises at the trailing edge and at the retracting leading edges. Images were acquired every 30 seconds and the movie is played at 30 frames/s. Scale bar is 10 μm.

